# Quantitative Characterization of Duodenal Gastrinoma Autofluorescence using Multi-photon Microscopy

**DOI:** 10.1101/2022.05.19.492747

**Authors:** Thomas G. Knapp, Suzann Duan, Juanita L. Merchant, Travis W. Sawyer

## Abstract

Duodenal gastrinomas (DGASTs) are neuroendocrine tumors that develop in the submucosa of the duodenum and produce the hormone gastrin. Surgical resection of DGASTs is complicated by the small size of these tumors and the tendency for them to develop diffusely in the duodenum. Endoscopic mucosal resection of DGASTS is an increasingly popular method for treating this disease due to its low complication rate but suffers from poor rates of pathologically negative margins. Multiphoton microscopy (MPM) is capable of capturing high-resolution images of biological tissue with contrast generated from endogenous fluorescence (autofluorescence) through two-photon excited fluorescence (2PEF). Second harmonic generation (SHG) is another popular method of generating image contrast with MPM and is a light-scattering phenomenon that occurs predominantly from structures such as collagen in biological samples. Some molecules that contribute to autofluorescence change in abundance from processes related to the cancer disease process (e.g., metabolic changes, oxidative stress, angiogenesis). MPM was used to image 12 separate patient samples of formalin-fixed and paraffinized DGAST slides with a SHG channel 4 2PEF channels, each tuned to capture fluorescence from NADH, FAD, lipofuscin, and porphyrin. We found that there was a significant difference in the relative abundance of signal generated in the 2PEF in comparison to the neighboring tissues of the duodenum. Texture extraction was used to create linear discriminant classifiers for tumor vs all other tissue classes before and after principal component analysis (PCA) of the texture feature dataset. PCA improved the classifier accuracy and reduced the number of features required to achieve maximum accuracy of the classifier. The LDA classifier after PCA distinguished between tumor and other tissue types with an accuracy of 90.6 - 93.8%. These results suggest that MPM 2PEF and SHG imaging is a promising label-free method for discriminating between DGAST tumors and normal duodenal tissue which has implications for future applications of in vivo assessment of resection margins with endoscopic MPM.

## 1. Introduction

### 1.1 Gastroenteropancreatic Neuroendocrine Tumors

Neuroendocrine tumors (NETs) are neoplasms that are thought to originate from secretory cells of the endocrine system and are characterized by their ability to express peptide hormones and biogenic amines. Highly heterogeneous in nature, NETs present with a wide spectrum of clinical features dependent on their primary site and molecular features resulting from their biologic, genetic, and epigenetic differences.^1^ The gastroenteropancreatic tract is a common site for NET formation and encompasses a wide variety of neoplasms that can be categorized based on their site of origin and secretory properties. Approximately 80% of gastroenteropancreatic NETs (GEP-NETs) are found in the small and large intestines, with a majority of these arising in the former.^2^ Small intestinal NETs account for ~2% of all gastrointestinal neoplasms and carry a higher potential for malignancy.^1^ SI-NETs are classified as ‘functional’ or ‘non-functional’ based on their hormone-secreting properties, with secretory tumors presenting in 30-40% of cases. Functional NETs of the small intestine predominantly secrete serotonin, somatostatin, or gastrin.^3^

Gastrin-secreting NETs, or gastrinomas, are a relatively rare group of GEP-NETs with an annual incidence of 0.05 – 0.2 cases per 100,000 ^4^ that develop in the pancreas and duodenum with 60 – 95% of cases occurring in the duodenum.^5^ Hypergastrinemia resulting from gastrinoma development may stimulate hyperchlorhydria due to the stimulatory effects of gastrin on gastric parietal cells.^6, 7^ The clinical manifestation of chronic gastric acid hypersecretion secondary to gastrinomas was first recorded by Zollinger and Ellison and is hence termed Zollinger-Ellison Syndrome (ZES).^8^ ZES typically presents as pain secondary to heartburn and/or peptic ulcer disease, diarrhea, abnormal weight loss, gastrointestinal bleeding, strictures, or perforations/penetrations of the GI tract.^9^ Hypergastrinemia from gastrinoma tumors has also been linked to the development of type I and type II gastric NETs^10^ which can regress following gastrinoma resection.^11^ Trends in NET cases show that this disease has been steadily growing in incidence over the last five decades which has been partially attributed to increased ‘awareness’ of the disease through research and improvements in screening and diagnostic technology.^3^

While conventional cross-sectional CT, MRI or ultrasound are commonly used for pre-operative detection and localization of gastrinomas, these modalities are at risk for missing duodenal gastrinomas (DGASTs) which tend to be small (<0.5cm) and develop diffusely, perpetuating the risk of future metastasis.^12^ Improvements in functional imaging using ^68^Ga-DOTATATE PET/CT has shown its promise in diagnosing and localizing primary tumor sites and areas of metastases due to its high sensitivity and specificity for primary and metastatic GEP-NETs using radiolabeled somatostatin receptor analogues.^13^ Endoscopic ultrasound (EUS) has been used to determine the depth and size of duodenal tumor growth which informs the surgeon whether an open surgical approach or endoscopic resection is appropriate.^15^ These modalities are both hindered in their ability to assess small lesions due to their low resolution, while EUS is highly operator dependent and can struggle to generate contrast in smaller lesions.^46^

The 2017 guidelines on the surgical resection of small bowel NETs from the North American Neuroendocrine Tumor Society recommend palpation of the full small bowel to ensure complete oncological resection. These guidelines extend towards the recommendation of resecting the small bowel through an incision or more invasive open abdominal surgery to facilitate adequate palpation.^14^ Less invasive surgical approaches such as endoscopic resection of duodenal NETs has gained popularity and is recommended for duodenal lesions < 10 mm in size by the European Neuroendocrine Tumor Society.^16^ While retrospective studies of endoscopic resection show this procedure as having relatively low complication rates, this method suffers from poor margin definition and a greater rate of recurrence in comparison to open surgery.^17^ Due to the drastic improvement in prognosis with full resection of these tumors,^5^ it is clear that methods for performing *in vivo* assessment of tumor clearance would greatly benefit patients undergoing this procedure.

### 1.2 Autofluorescent Multi-photon Microscopy

The detection of endogenous fluorescence, or autofluorescence (AF), has shown promise in its use as an ‘optical biopsy’ tool, or a means of distinguishing between tissue types and healthy versus diseased tissue based on their innate fluorescent properties. ^18–21^ AF imaging provides a minimally invasive method of probing tissue characteristics without the use of exogenous contrast agents, eliminating the risk of toxicity or potential reaction to the dyes. A major limitation of AF imaging is the relatively low signal-to-noise ratio produced when sensing light emitted from the naturally occurring fluorescent molecules within tissue which can lead to a high false-positive rate.^22^ This can be mitigated with careful characterization of the AF properties of diseased and normal tissue. For example, quantification of AF has been shown to be a promising method of distinguishing between adenomatous and hyperplastic colon polyps^19^ and normal, hyperplastic, and dysplastic cells of isolated colon crypts.^20^ Multi-photon microscopy (MPM) is a promising modality for quantifying tissue AF at a high resolution and contrast. Two-photon excited fluorescence (2PEF) is one method of generating image contrast in MPM and is the process of an endogenous fluorophore absorbing two photons within a narrow enough time interval to create fluorescence emission. The two-photon absorption process utilizes light of longer wavelengths (e.g., 700 - 1000 nm) to excite fluorescing electrons to a similar degree as single-photon fluorescence, which uses light of a shorter wavelength, (e.g., 360 – 550 nm), i.e., a greater photon energy. Second-harmonic generation (SHG) is another popular method of creating MPM contrast and is a nonlinear light scattering phenomenon from non-centrosymmetric structures, which is predominantly collagen in tissue samples.

Use of excitation light of longer wavelengths in MPM allows for imaging deeper into tissue samples due to the inverse relationship between wavelength and light scattering as it passes through a medium. This would play an essential role for probing through tissue layers deep within the layers of the small intestine.^23^ Signal generation from 2PEF and SHG events scales with the square of the excitation irradiance, confining photon emission from the tissue within the point of focus. The confined space of these events results in excitation of fluorophores within the volume of tissue from which the MPM optics can fully focus onto the detector. This benefits in the imaging of biologic samples by reducing the chance of inadvertent tissue damage or photobleaching during image acquisition and fluorescence emission outside of the focal plane which could degrade image quality.^25^ Ultimately, MPM represents an imaging modality that is capable of acquiring volumetric images at a sufficiently high resolution to enable the accurate assessment of the molecular features comprising a tissue specimen. Developing models of DGAST multiphoton fluorescence is an initial step towards *in vivo* label-free measurements of these lesions. This would bypass the need for fluorescent dyes that pose the risk of cytotoxicity while allowing for screening, diagnosis, and staging of DGASTs.

### 1.3 Multiphoton Image Analysis

Use of a tunable laser light source and multi-channel detector array in MPM of tissue AF allows for the selection of excitation and emission wavebands that are more likely to collect signal from different proportions of fluorescent molecules. For instance, imaging of fluorescent species such as NADH and FAD that are linked to established abnormalities in tumor cells, e.g., the Warburg effect^26^ creates a contrast between regions of tumor and surrounding tissue. Other fluorophores with relevance to tissue changes during the cancer disease process include lipofuscin, porphyrins, riboflavins, collagen, elastin, and amino acids.^47^

Quantification of tissue AF and differences in the AF between tumor and normal tissue can be accomplished in several ways. A straightforward approach is the thresholding and direct comparisons of 2PEF and SHG signal abundances.^27^ While east to interpret, this approach is strongly influenced by variations in light source power, instrument alignment, and tissue preparation. A more robust approach towards quantification is texture analysis, which can provide the spatially-varying statistical characteristics of image tone and texture through the co-occurrence of pixel gray-level values.^28^ Texture analysis is valuable to characterize the distribution of image brightness, which can describe features such as contrast, homogeneity, and entropy within the image. In general, texture analysis has been used with success in the context of tissue characterization for cancer imaging, including ovarian cancer using optical coherence tomography^21^, breast cancer lesions with MRI^48^, and non-small cell lung cancer using non-contrast-enhanced CT.^49^ Several different methods of texture analysis exist, which can be classified into four broad categories of statistical methods, structural, model-based, and transform-based methods. Each approach can provide different information about the image content and how that information is contained within the spatial distribution of pixel / voxel values.^50^ This information can then be used to train classifiers to distinguish between different classes within images, which in this case would be tissue types.

In this work, we aim to assess the suitability of MPM for tissue characterization and margin definition of DNETs. To our knowledge, this is the first attempt at optically characterizing DGAST tumors with MPM. We collect a set of 2PEF and SHG images of fixed human DGAST samples from 12 patients. Image regions of interest (ROIs) corresponding to tumor tissue and surrounding regions of duodenal tissue are quantified by calculating the magnitude of fluorescence emission, as well as conducting statistics-based gray-level co-occurrence matrix (GLCM) texture analysis. Our results show that multi-photon imaging provides excellent label-free contrast for identifying and delineating cancerous tissue. We find that statistically significant differences in AF and texture features between DGASTs and normal tissue of the duodenum exists. We then developed classifiers using the texture feature and show that the tumor tissue can be successfully differentiated from surrounding tissues with over 90% accuracy. These results are promising and demonstrate that MPM AF imaging can be used as a tool for demarcating areas of DGASTs from healthy tissue of the duodenum. Use of MPM in this context can eventually fill a vital role in the process of screening, localizing, and assessing surgical margins when combined with endoscopy or laparoscopy.

## 2. METHODS

### 2.1 Sample Acquisition and Preparation

Formalin-fixed paraffin-embedded (FFPE) slides of GEP-NETs were obtained from the University of Michigan Endocrine Oncology Repository (IRB #HUM00115310) through a Materials Transfer Agreement between Dr. Tobias Else and Dr. Juanita Merchant. Original tissue specimens were collected from human subjects who received a diagnosis of Multiple Endocrine Neoplasia (MEN1) syndrome, gastrinoma, or both. Slides containing tissue that was not confirmed gastrinoma, either through positive staining or hypergastrinemia pre-surgery were excluded from this study. Informed consent was given by the patient prior to sample collection and all specimens were de-identified to protect patient confidentiality. Samples were collected during upper endoscopy or during surgical procedure and placed in 10% neutral-buffered formalin prior to paraffin embedding and sectioning. Five-micron FFPE sections were used for downstream imaging studies. In total, 12 separate slides, some including multiple histological sections (19 sections total) were used for this study.

### 2.2 Multiphoton Imaging

Specimens were imaged using a Zeiss LSM 880 microscope (Carl Zeiss Microscopy, White Plains, NY) with a tunable MaiTai HP titanium:sapphire light source (Spectra-Physics, Mountain View, CA). A 690+ nm long-pass dichroic mirror and a 660nm short-pass emission filter were used as the main beam splitter and secondary beam splitter in the invisible light beam path. Coverslips were dry-mounted onto the slides and images were taken at a 700 – 800 master gain and 50 mW average laser power for each of the fluorescence channels and the SHG channel using a plan-apochromat 20X/0.8 NA objective (420650-9902-000, Carl Zeiss Microscopy, White Plains, NY). Gain was adjusted on a per-sample basis and held fixed while acquiring the separate imaging channels to allow for the direct comparison of signal intensity from the separate tissue sites within the image. In total, five separate image channels were collected. The 2PEF excitation and emission bands (**Table 1**) were determined using the known two-photon fluorescent spectra of NADH^29,30^, FAD^30^, porphyrin^31^, and lipofuscin.^32^ We selected these fluorophores as they are established biomarkers of cancer development (e.g., through irregularities in cell metabolism, senescence, and vascularization) and have well-characterized fluorescent properties.

**Table 1:**
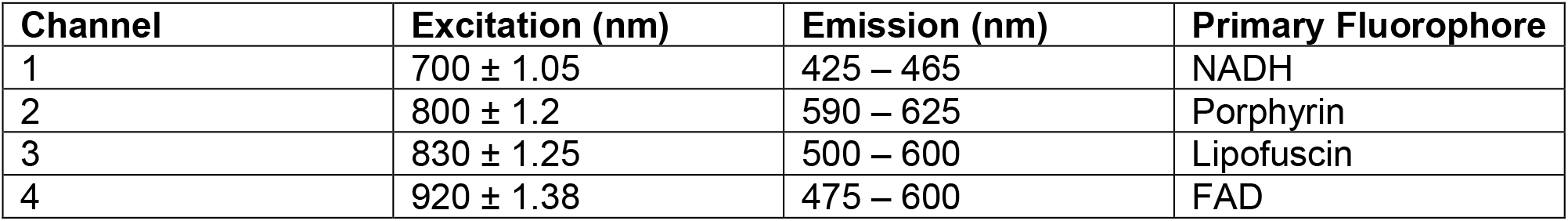
Excitation and emission (detection) bands for the 4 2PEF imaging channels alongside the fluorophore we expect to be producing the dominant signal of the channel.

| Channel | Excitation (nm) | Emission (nm) | Primary Fluorophore |
| --- | --- | --- | --- |
| 1 | 700 ± 1.05 | 425 – 465 | NADH |
| 2 | 800 ± 1.2 | 590 – 625 | Porphyrin |
| 3 | 830 ± 1.25 | 500 – 600 | Lipofuscin |
| 4 | 920 ± 1.38 | 475 – 600 | FAD |

Large area scans were acquired by collecting individual image tiles of 256 x 256 pixels in size. Images were obtained with 16-bit gray scale resolution and 10% overlap between the separate tiles using an automatic motorized stage. Total area covered by the tiling scan was selected depending on the amount of each tissue type captured within the scan area for adequate ROI sampling within the image and analysis of bulk tissue morphology via image feature extraction. Our criteria for selecting the image size were based on the density of tissue types within the scan area, such that there were no artifacts and less than 90% brightness variation within the area to draw at least three 0.25 mm^2^ ROIs for each class contained in the image. The tile-scanned images were collected in z-stacks to address drop-off in brightness that occurs over the uneven surface of the sample secondary to the sectioning process. The number of acquisitions in the z-stack was selected depending on the site and severity of brightness variations. Z-stacks were acquired such that over 90% of the sample area was in-focus with the image, ranging between 3-9 acquisitions in the z-axis.

### 2.2. Image processing

To process the acquired images, a flat-field correction was first applied to account for illumination variations. This was accomplished by averaging the pixel brightness for each 256 x 256 tile over all tiles and fitting the result to a linear brightness trend. All collected tiles were then divided by this trend to account for the illumination variation in the laser irradiance. The tiles were then combined using the ImageJ Grid/Collection stitching plugin^33^ and a maximum projection image was created using the Z-stack images for the separate image channels. A custom MATLAB frequency filtering step was then used on the maximum projection images. For a more complete writeup on this image processing, please refer to Knapp et al.^34^ A MATLAB implementation of the structural similarity index (SSIM)^35^ was used to determine if image information was retained post-processing. Visual examination of the images was done to confirm the absence of artifacts and the SSIM score was used to quantitatively support the inclusion of the images in our analysis. An example of the processing steps and assessment with SSIM is shown in **Figure 1**.

**Figure 1:**
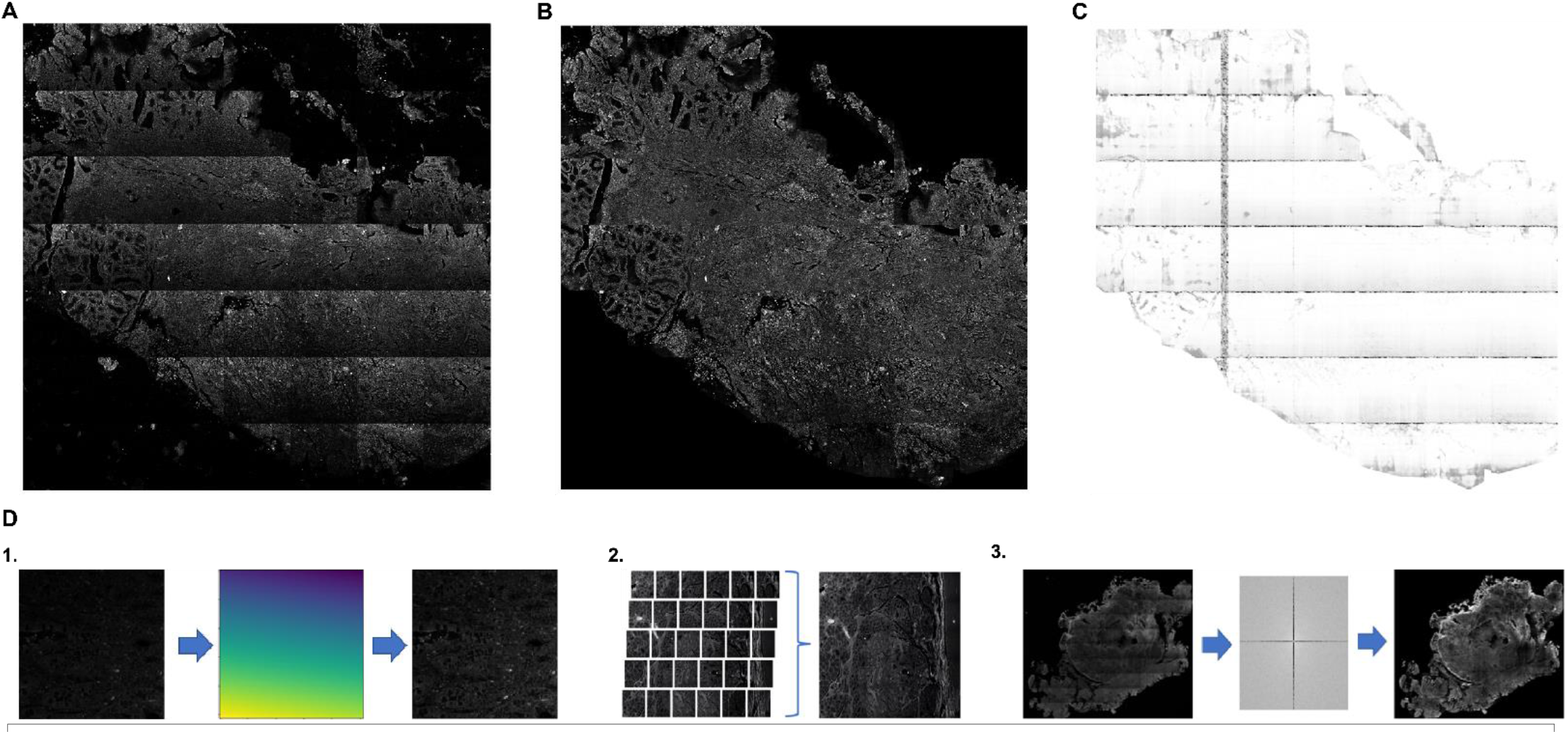
2PEF channel pre-processing (A) and post-processing (B) showing the presence and removal of the tiling artifact. The SSIM image of the same region post-processing (C) shows good retention of image structural information, indicated by the peak value of 1 represented as white pixel values in areas where image structure was retained through the processing steps. The dark lines on the SSIM show where the gridline artifact was removed. On the bottom row of the figure (D) is an abbreviated illustration of the image processing steps. 1 = custom flat-field approach for correcting image tile brightness variations, 2 = ImageJ stitching methods to combine image tiles, 3 = frequency filtering to remove residual gride lines following the combination of the image tiles.

ROIs from the 2PEF and SHG imaging channels were extracted using ImageJ software^36^ for the separate tissue types present in the image, classified as either stroma, villi / lamina propria, Brunner’s glands, abnormal Brunner’s glands, and tumor. ROIs were individually inspected, and any containing artifacts were excluded resulting in a total of 209 ROIs included in our analysis. Co-author JLM performed ground-truth classification of tissue using morphologic information and immunohistochemistry (IHC) stains of serial sections of our samples as shown in **Figure 2.**

**Figure 2:**
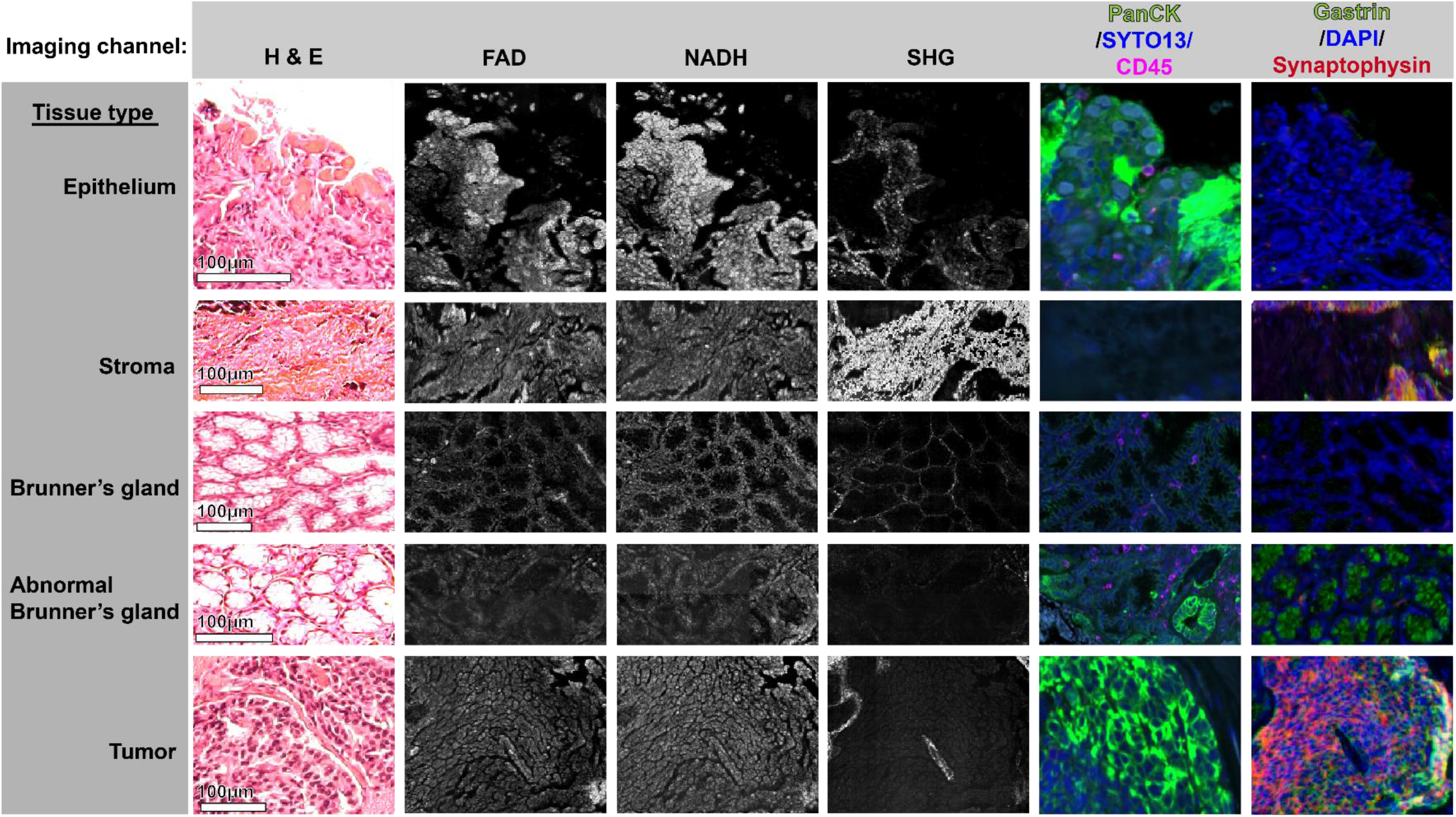
ROIs of the different tissue types showing examples of differences in morphology and IHC staining used for classification. The regions of gastrinoma tumor were verified through gastrin and synaptophysin staining. Regions of collagenous stroma produced a strong SHG signal with a distinct fibrous morphology. Villi and lamina propria were classified together and determined to be regions of the sample obviously oriented towards the lumen in-situ and/or strongly stained by the epithelial marker PanCK and having the characteristic appearance of villi. Brunner’s glands had a characteristic glandular appearance in 2PEF, SHG,H&E and IHC stained I,ages, while abnormal Brunner’s glands had an atypical expression of epithelial markers that shown with PanCK staining.

Synaptophysin and chromogranin A are traditional markers for NETs. As gastrin-expressing NETs have been shown to stain weakly for chromogranin A^51^, synaptophysin was used in combination with a gastrin stain to validate that the tumor regions were gastrin-expressing NETs, i.e., gastrinomas. PanCK is an epithelium marker used for determining tissue morphology during staining and has been used for identifying tumor-associated, or abnormal, Brunner’s glands.^52^

### 2.3 Quantification and Comparison of 2PEF and SHG

All analysis was completed using Python 3. A threshold was first applied to each image to remove saturated pixels and to limit the influence of background signal, as well as the inclusion of pixel values containing noise. Lower thresholds were determined for each channel by taking the average value of an image area with no tissue to determine background (**Table 2**). For the FAD-dominant channel, there were strong contributions in fluorescence from lipofuscin deposits that manifest as localized bright spots, which is a common observation when measuring this biomarker.^53^ This channel was set with an upper threshold of 50000 to remove the influence of these features (**Table 2, channel 4)**. Following thresholding, the mean 16-bit pixel value was recorded for each ROI and the measured values were compared between all tissue types within each image. Comparison of fluorescent and SHG intensity was constrained to within-image sample ROIs (e.g., paired comparisons between ROIs for each patient) to eliminate variations between the tissue slide sectioning, and instrument-level variations between image acquisition sessions. As the ROIs only differed in their optical contrast, a paired t-test was used to determine the degree of significance between the comparisons of relative 2PEF and SHG signal intensity between the tissue classes. For all statistical comparisons, we averaged the values of all ROIs created for each tissue class within the slides from each patient to be treated as a different sample (n = 12).

**Table 2:** Values of the global thresholds that applied to the 2PEF and SHG image channels. Channel 4 was the FAD-dominant excitation/emission band. Due to the broad detection band, a high proportion of bright lipofuscin deposits were present in the images of this channel, hence the lowered upper threshold. Lower thresholds were based on averaging of the image background noise.

| Channel | Lower Threshold (16-bit pixel value) | Upper Threshold |
| --- | --- | --- |
| 1 | 3000 | 65535 |
| 2 | 3000 | 65535 |
| 3 | 3000 | 65535 |
| 4 | 1500 | 50000 |
| SHG | 1000 | 65535 |

### 2.4 Texture Analysis and Classification

We applied texture analysis to the collected ROIs from each of the MPM imaging channels. Haralick’s method of extraction^28^ based on the GLCM was used to generate thirteen values describing the texture of each ROI. Prior to texture extraction, the ROIs were normalized and reduced to an 8-bit data range. The GLCM describes the spatial distribution of pixel gray-level values within an image. Values of the GLCM p(*i, j* | *d, θ*) are the measured occurrences of a gray-level (*i*) being a distance, *d*, away from gray-level (*j*) in the direction *θ*, where *θ* is 0 deg, 45 deg, 90 deg, and 135 degrees for a two-dimensional image. The thirteen texture values calculated from the GLCM were averaged for the four directions of *θ*. With five MPM channels, this results in a total of 65 features for each ROI.

1. *Angular Second Moment*:

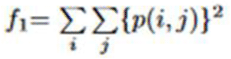
2. *Contrast*:

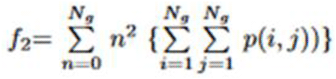
3. *Correlation*:

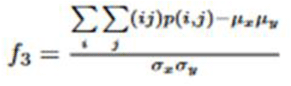
4. *Sum of Squares*: *Variance*

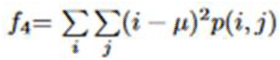
5. *Inverse Difference Moment*:

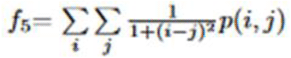
6. *Sum Average*:

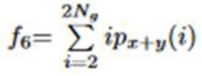
7. *Sum Variance*:

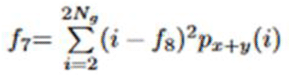
8. *Sum Entropy*:

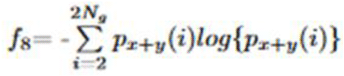
9. *Sum Entropy*:

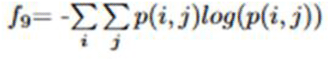
10. *Dif ference Variance*:

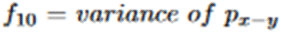
11. *Difference Entropy*:

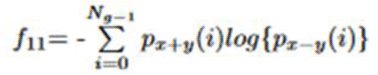
12. *Information Measures of Correlation*:

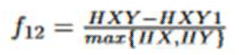
13. 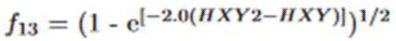

Equations defining the texture features extractable from the GLCM of images.

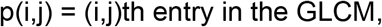

p_x_(i) = *i*th entry in the marginal-probability matrix created by summing rows of 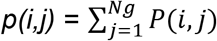,

N_g_ = number of distinct gray levels in quantized image,

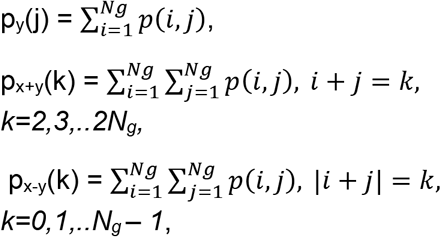

μ_x_, μ_y_, σ_x_, σ_y_ are the means and standard deviations of p_x_ and p_y_.

The ability to classify the collected ROIs was tested using different combinations of the subsets of the 65 texture features in Linear Discriminant Analysis (LDA) model using singular value decomposition.^37^ We examined the classification performance between tumor and the four tissue types of villi / lamina propria (grouped as epithelium), stroma, Brunner’s gland (BGs), and abnormal Brunner’s gland (aBGs) to assess the quantitative ability to differentiate tissue images. The performance of the classifier to distinguish between normal and abnormal BGs was also assessed, as the occurrence of aBGs could represent a transition from normal tissue to disease state (e.g., pre-cancerous tissue). We first test the LDA classifier using an input of the raw texture feature values and then evaluated the model after applying Principal Component Analysis (PCA) for dimensionality reduction of the feature space. PCA was applied to the dataset containing all ROIs and features, and the top 15 principal components were retained while the others discarded. The top 15 components accounted for 97% of the variance in the dataset.

We then compared the classification performance of different sizes of feature families from this dataset (i.e., families composed of different numbers of features). We tested feature families consisting of subsets ranging from two features to ten features, as texture classifiers have been shown to plateau in performance at around 6 or fewer features.^21^,^39^,^40^ This process was repeated for the raw feature values, although it required considerably longer computation time to iterate through each feature combination.

We used a leave-one-out approach to measure the accuracy of classifiers using the feature families. We performed an exhaustive comparison of all possible feature families for each size. The top performing feature subsets for each feature subset family were selected as optimal for each size, and the classification accuracy (defined as the mean score of the classifier correctly separating members into their own class) and receive operator characteristic were recorded.

## 3. RESULTS & DISCUSSION

### 3.1 Qualitative Assessment of Tissue Contrast from 2PEF and SHG

Clear morphologic differences between the separate tissue classes can be distinguished with qualitative assessment of the 2PEF and SHG images. **Figure 3** shows the different morphologic information provided by SHG (**Figure 3** B) and 2PEF (**Figure 3 C-F)** imaging. There is a clear increase in cell density within the tumor regions shown in the 2PEF images while SHG shows the disorganized and thick stromal (S) changes near the tumor sites which is consistent with prior findings of desmoplastic changes occurring around tumor regions.^41^ There is a loss of SHG signal and the appearance of thick collagenous encapsulation of the neoplasms. In comparison, the 2PEF and SHG images of the neighboring Brunner’s glands (BGs) show clear contrast to the tumor in the organized arrangement of the glands with thin collagen walls composing their structure. Immunostaining of the DGAST samples (**Figure 3, G)** shows a high level of gastrin and synaptophysin expression within the tumor regions which is characteristic of DGASTs.^45^ While clear distinctions between normal and abnormal BGs (aBGs) could not be made with qualitative assessment of 2PEF and SHG imaging, aBGs were shown to stain for gastrin and the epithelial marker PanCK. aBGs were typically located near tumor regions and were demarcated due to BGs being a suspected precursor to DGASTs.^42,52^ H & E staining of the DGAST samples (**Figure 3, B 3)** shows the trabecular patterning of DGASTs^43^ in the tumor region corresponding to the marked (T) region of the NADH channel image.

**Figure 3:**
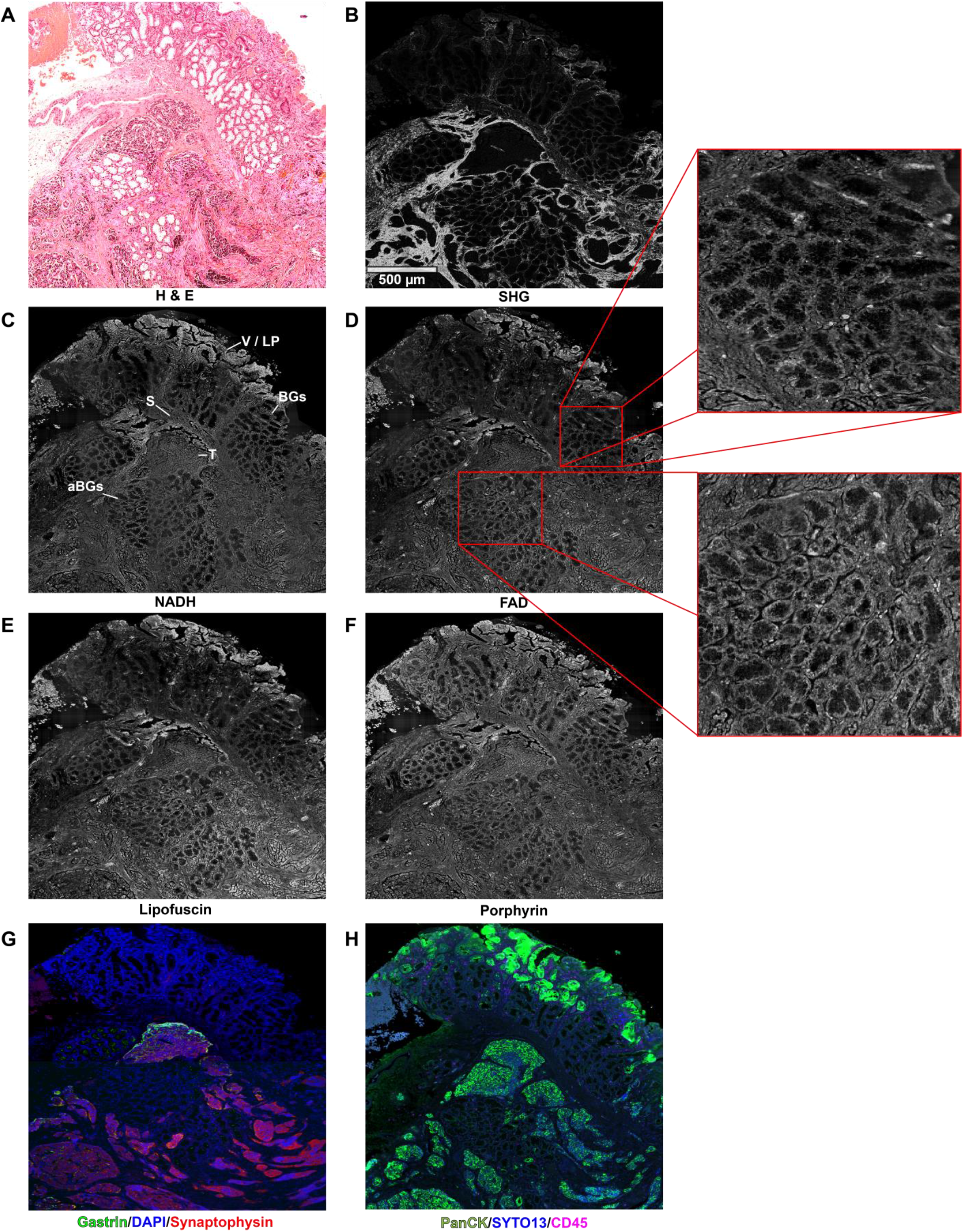
H & E (A), SHG (B), 2PEF channels (C,D,E,F), and IHC-stained images (G,H) of a duodenal gastrinoma specimen. Tissue classes are labeled in the NADH-dominant channel image (C), V / LP = villi /lamina propria, BGs = Brunner’s glands, S = stroma, aBGs = abnormal Brunner’s glands, T = tumor. Gastrin staining (G) is specific for gastrin-expressing cells, while synaptophysin is specific for neuroendocrine tumors. PanCK stain (H) is specific for epithelial markers, SYTO stains DNA, and CD45 for lymphocytes (immune cells). aBGs were classified based on their abnormal expression of epithelial marker and gastrin which was determined with PanCK and gastrin staining. Zoomed views show BGs and aBGs imaged with the FAD-dominant 2PEF channel. The H & E images of the DGAST samples were used to assess tumor morphology and as landmarks for regions of tumor neighboring other tissue types of interest for comparing the measurable autofluorescence.

**Figures 4–6** show H&E, SHG, and 2PEF images for three other patient samples. In **Figure 4**, we see a comparison of bordering BGs and aBGs near a large region of tumor. While not yet clear in its significance, there appears to be a reduction in the SHG signal (**Figure 4 B**) and thickening of the gland tissue seen in the FAD-dominant 2PEF channel (**Figure 5 C**) seen in the aBGs. **Figure 5** shows the bordering region of a tumor and normal BGs. **Figure 5 B** shows the loss of SHG signal within the tumor that is normally occurring within the gland structure. The tumor exhibits many bright deposits in the lipofuscin-dominant 2PEF channel (**Figure 5 C**). **Figure 6** is an example of the thickened stroma surrounding sites of tumor which is accentuated in SHG imaging (**Figure 6 B**). Many bright deposits were present in the DGAST region when imaged with the porphyrin (**Figure 6 C**), lipofuscin, and FAD-dominant channels. These clusters are thought to be dye used for tumor localization prior to resection and/or accumulation of lipofuscin deposits.

**Figure 4:**
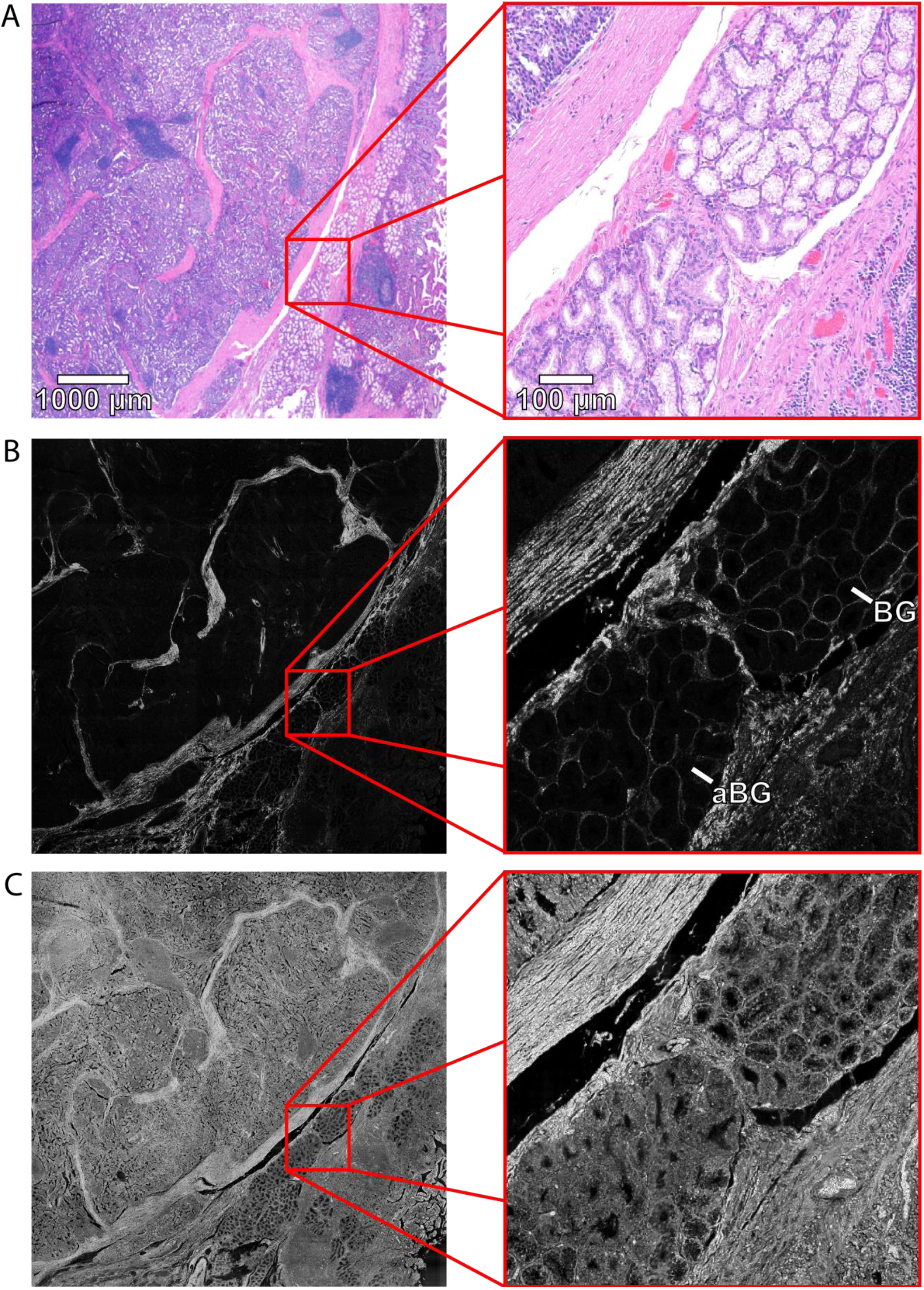
H&E (A), SHG (B) and FAD-dominant (C) channel images of DGAST tumor with zoomed views showing the border between abnormal (aBG) and normal Brunner’s glands (BG).

**Figure 5:**
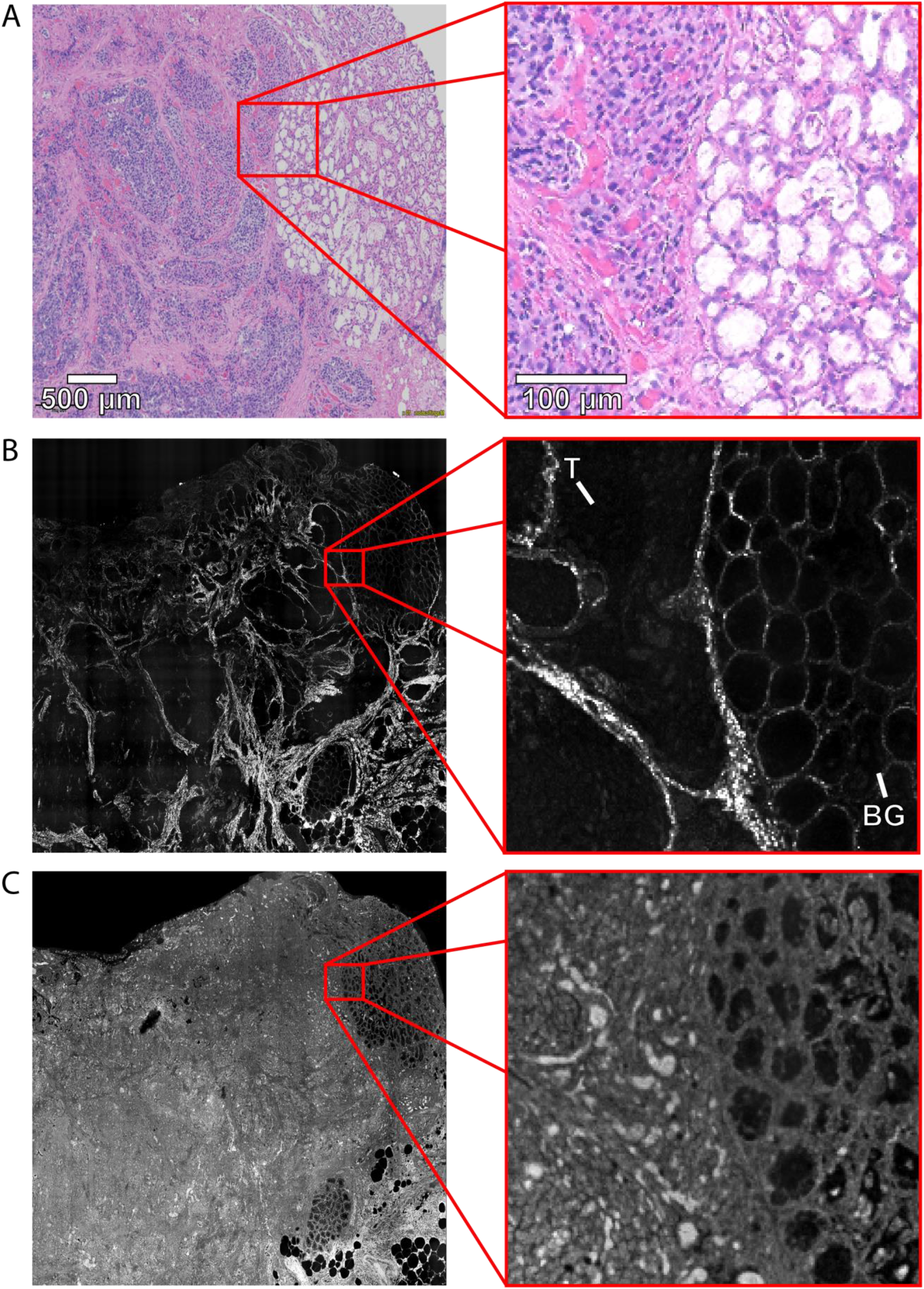
H&E (A), SHG (B) and lipofuscin-dominant (C) channel images of DGAST tumor with zoomed views showing the border between tumor (T) and Brunner’s glands (BG).

**Figure 6:**
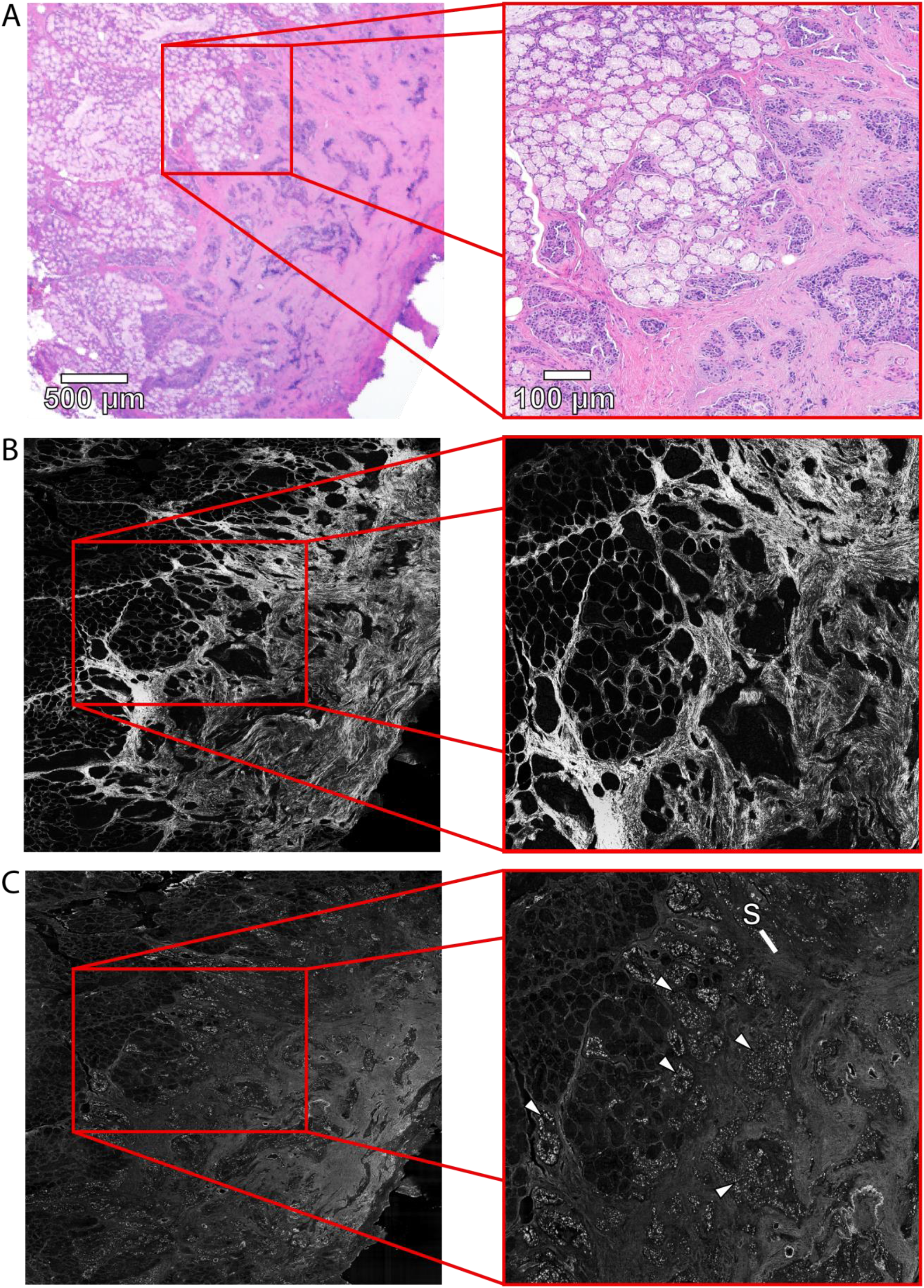
H&E (A), SHG (B) and porphyrin-dominant (C) channel images of DGAST tumor with zoomed view showing the highly disorganized stroma (S) with the characteristic trabecular appearance surrounding the diffuse clusters of tumor (marked with arrows in C).

### 3.2 Quantitative Assessment of 2PEF and SHG Image Channel Intensities

T-test comparisons of the raw relative mean signal magnitude collected from the tissue classes in the separate 2PEF and SHG imaging channels (**Figure 6)** shows a significant difference (p < 0.001 for the NADH, lipofuscin, and porphyrin-dominant channels, (**Figure 6 A, C, D**), p < 0.01 for the FAD-dominant channel (**Figure 6 B**)) in 2PEF between the tumor regions and glandular tissue. Epithelial tissue (villi and lamina propria) exhibited a significantly greater signal in the lipofuscin (p < 0.05, **Figure 6 C**) and NADH-dominant (p < 0.001, **Figure 6 A**) channels in comparison to the tumor tissue. This reflects earlier findings of broad-spectrum AF signal emitted predominantly from lipofuscin and lipofuscin like pigment within the lamina propria of mice and humans.^53^ As expected, stromal tissue produced a significantly greater (p < 0.001, **Figure 7 E**) SHG signal in comparison to the epithelial, glandular, and tumor tissue classes.

**Figure 7:**
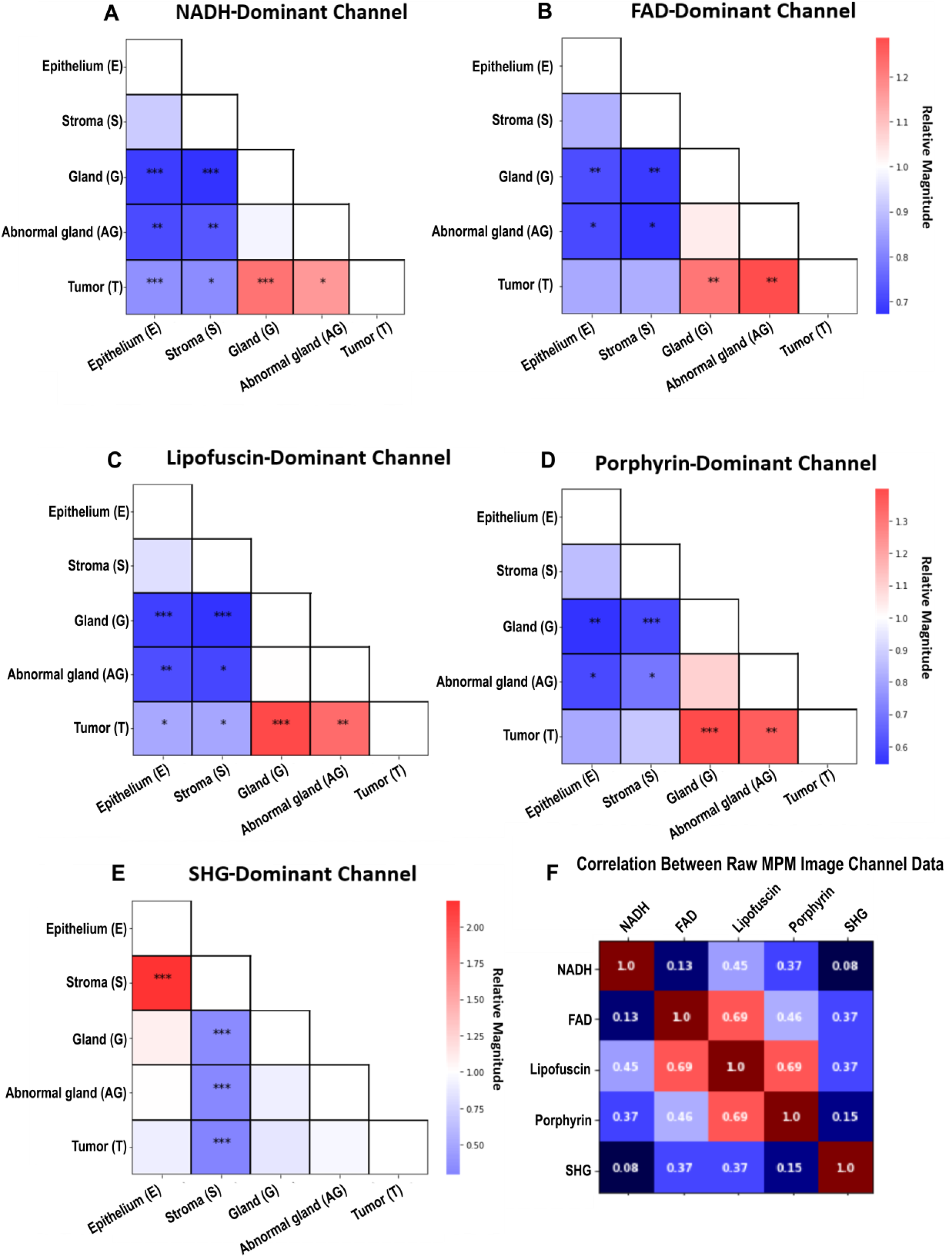
Heat matrices of the relative raw signal intensity comparisons between each tissue class. * = p < 0.05, ** = p < 0.01, *** = p <0.001. The matrices are interpreted as the row divided by the column. The correlation matrix (F) shows a high positive correlation between the lipofuscin, FAD, and porphyrin-dominant image channels prior to normalization.

A correlation matrix of the image channel data was produced to assess crosstalk between imaging channels (**Figure 6 F)**, which suggested that a large degree of overlap existed between the lipofuscin, FAD, and porphyrin channels. This was unsurprising considering the broad emission band of lipofuscin^44^ and comparisons were made following normalization of each ROI through division of the corresponding lipofuscin-dominant image channel as shown in **Figure 8**. The correlation matrix of the normalized data (**Figure 7 F**) shows a lower degree of overlap between the image channels. There was a mild decrease in the level of difference of the 2PEF signal magnitudes between the tissue types. Notably, the tumor tissue no longer produced a significantly greater magnitude of signal in the NADH and porphyrin-dominant channels in comparison to the aBGs tissue class. While 2PEF signal magnitude decreased in all channels following normalization, the greater loss in signal for the abnormal glands may suggest that lipofuscin accumulation plays a larger role in the AF signal produced by this tissue in comparison to normal Brunner’s glands.

**Figure 8:**
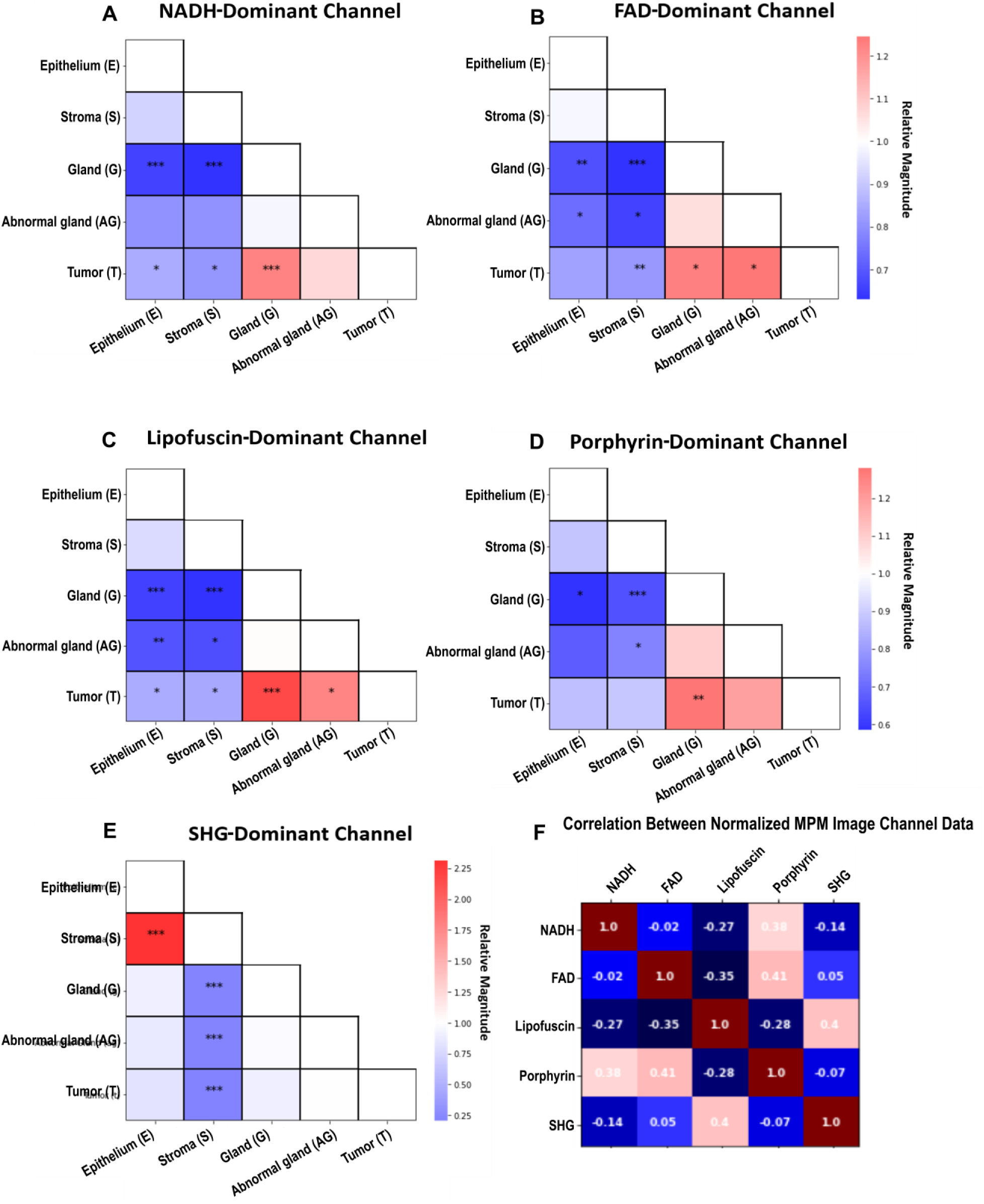
Heat matrices of the relative signal intensity comparison between tissue classes post-normalization by the lipofuscin imaging channel. * = p < 0.05, ** = p < 0.01, *** = p <0.001. The correlation matrix (F) shows a lower degree of correlation between the lipofuscin, FAD, and porphyrin channels as expected. This does result in a decrease in the deg**r**ee of di**ff**erence between the tissue classes for the FAD and porphyrin-dominant channels. A significant difference between the tumor and BGs still exists for the porphyrin-dominant channel and the tumor, stroma, and normal/abnormal glands for the FAD-dominant channel.

**Figure 9:**
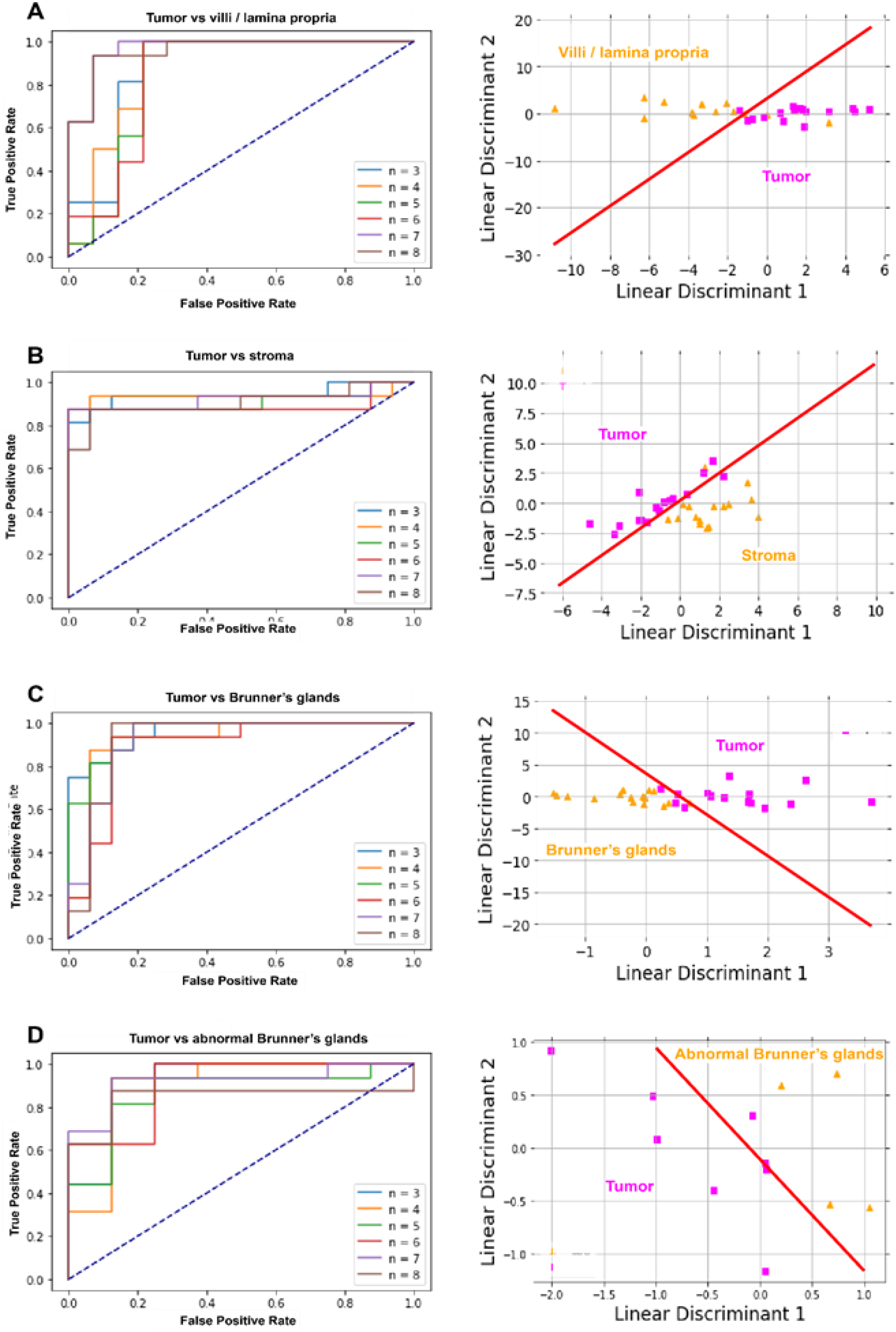
Receiver operating characteristic curves of the LDA classifier and LDA projections with hyperplane plotted in red distinguishing between tumor and BGs, aBGs, VLP, and stroma. n = number of features used to generate classifier. LDA projections for the tumor vs BG, stroma, and VLP classifier are shown with classification using 6 features, the tumor vs aBG plot is with 4 features used for classification.

### 3.2 Classification Using Texture Features

**Table 3** shows the results of the LDA accuracy for classifying the tumor vs all other tissue classes and BGs vs aBGs using the extracted texture features from the ROIs prior to PCA. We chose these comparisons due to the clinical relevance of tumor localization and the identification of potentially pre-cancerous tissues as described by the aBG state. The results show that, using raw feature values, the accuracy for classifying tissue types tends to plateau with five to six features. The accuracy for delineating tumor, BG, aBG, and VLP all exceeds 90%, yet is limited to 71.9% for stroma. Additionally, we see that the accuracy for classifying BG and aBG is maximized at 79.2%. For most comparisons, we see a decrease in accuracy with the addition of features after the plateau point, which is a common observation when performing feature selection and classification with high-dimensional datasets. This plateau is reached when the additional features no longer increase the information content for classification. This observation suggests that six features can capture most of the relevant information.

**Table 3:** Number of features prior to principal component analysis included in the classifier and the resulting mean level of accuracy of the classifier in separating the tissue groups.

| Number of features used in classifier | % accuracy for classifying BGs vs aBGs | % accuracy for classifying tumor vs BGs | % accuracy for classifying tumor vs aBGs | % accuracy for classifying tumor vs VLP | % accuracy for classifying tumor vs stroma |
| --- | --- | --- | --- | --- | --- |
| 2 | 75.0 | 87.5 | 83.3 | 93.3 | 62.5 |
| 3 | 75.0 | 87.5 | 87.5 | 96.7 | 65.6 |
| 4 | 75.0 | 87.5 | 91.7 | 96.7 | 68.8 |
| 5 | 79.2 | 90.6 | 91.7 | 96.7 | 68.8 |
| 6 | 75.0 | 90.6 | 91.7 | 96.7 | 71.9 |
| 7 | 79.2 | 90.6 | 87.5 | 96.7 | 68.8 |
| 8 | 75.0 | 90.6 | 83.3 | 96.7 | 68.8 |

Principal component analysis (PCA) was used to reduce the dimensionality of the large set of texture features and the top 15 principal components were used in the LDA classifier. Results of the LDA accuracy for classifying tumor vs all other tissue classes using the texture features following PCA are shown in **Table 4**.

**Table 4:**
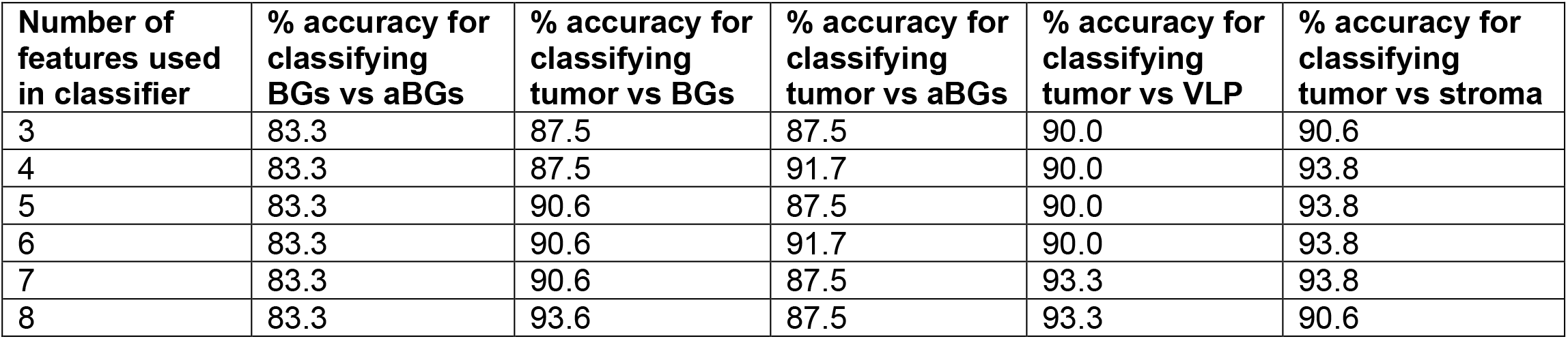
Number of features post-principal component analysis included in the classifier and the resulting mean level of accuracy of the classifier in separating the tissue groups.

Dimension reduction with PCA had an insignificant effect in improving classification accuracy between tumor and aBGs; however, a high level of accuracy was achieved with raw feature values. We do observe that, with PCA applied, the maximum accuracy is achieved including only two features. There was a marked increase in accuracy for the classification of tumor vs stroma following PCA (71.9 to 93.8%) and mild improvement in performance for classifying tumor vs BGs (90.6 to 93.8%) and BGs vs aBGs (79.2 to 83.3%). The classifier’s accuracy in separating the tumor and VLP classes worsened following PCA (96.7 to 93.3%). Overall, the maximum accuracy was also reach with fewer features overall. The decrease in accuracy for the classification of tumor and VLP is not surprising, as while PCA will identify the directions of maximum variance in data, it does not necessarily lead to the maximum separability between all groups. However, considering the marked increase in classification accuracy between the other four groups and the achievement of maximum accuracy with fewer features, the results suggest that PCA is an effective technique for dimension reduction and feature selection with this dataset.

Receiver operating characteristic (ROC) curves and LDA projections for tumor vs VLP, stroma, and BGs using the first two linear discriminants were generated with the PCA transformed texture feature data as shown in **Figure 8**. The ROC curves show a similar trend in the plateauing accuracy of classifiers using texture features^21,39,40^, although this plateau did range between 4-8 features depending on the tissue class.

## 4. CONCLUSION

In this work, we have shown that it is possible to distinguish between DGAST tumors and the normal duodenal tissue surrounding these lesions using multi-channel AF images collected with MPM. The separate tissue classes showed clear morphological differences in both the 2PEF and SHG image channels which corresponded well with H & E and immunostained images of the tissue. Collection of multiple 2PEF channels provided a robust means of differentiating between tissue classes through varying levels of significant differences existing in signal magnitude depending on the tissue and imaging channel. Imaging channels that collected signal dominated by fluorophores tied to cell metabolism and senescence, such as NADH/FAD and lipofuscin, respectively, provided a high degree of contrast between tumor and the neighboring BGs (p < 0.001, p < 0.05, p < 0.001 for each channel post-normalization). Normalization of the imaging channels showed that significant differences in 2PEF signal intensity exists between the DGAST tumors and normal tissue even while controlling for the high cellular density of the cancer. Texture features (e.g., contrast, entropy, autocorrelation) were extracted from the GLCM of ROIs and averaged from the five separate image channels for each tissue type. Subsets of the 65 texture features were used to create classifiers through single value decomposition linear discriminant analysis that classified the DGAST tumor vs BGs, aBGs, VLP, and stroma tissue with accuracies of 90.6%, 91.7%, 96.7%, and 71.9%, respectively and BGs vs aBGs with an accuracy of 79.2%. Principal component analysis (PCA) was used to reduce the dimensionality of the 65 texture features to 15 principal components containing 97% variance of the data. A new classifier using subsets of the transformed data showed improved classification accuracy between tumor vs BGs (90.6 to 93.8%), tumor vs stroma (71.9 to 93.8%), and BGs vs aBGs (79.2 to 83.3%). PCA transformation did mildly diminish the accuracy of the classifier for distinguishing between the tumor and VLP classes (96.7 to 93.8%). Use of PCA resulted in maximum accuracy with fewer features included in the subset of data used in creating the model for the BG vs aBGs, tumor vs aBGs, and tumor vs stroma classifiers. These early results show promising use of AF as a label-free approach for determining regions of DGAST tumors and potential for differentiating between areas of normal and abnormal glandular tissue which may be areas of early neoplastic change. With the high-resolution achievable with optical imaging, refinement of this classification process may offer the ability to perform margin analysis, screening, and/or diagnosis of duodenal NETs *in vivo*. Optimization of the classifier will be approached through further assessment of the amount of AF information required to produce discrete levels of accuracy (i.e., the optimum degree of information needed to collect during imaging) and how downsampling the images modifies classification using texture analysis. The combination of streamlining data collection and feature extraction while limiting the data size will make real-time analysis of duodenal NETs and the surrounding tissue during multi-photon imaging feasible.

